# An urbanized phantom tributary subsidizes river-riparian communities of mainstem gravel-bed river

**DOI:** 10.1101/2023.11.07.565958

**Authors:** JN Negishi, YY Song, I Matsubara, N Morisaki

## Abstract

Urbanization transforms natural river channels, and some rivers become invisible over time. How and whether the subsurface domains of the original waterways and aquifers connecting them (a phantom of historical landscape) are functional is not known. This study examined the effects of tributary groundwater (GW) inflow on the response of river-riparian organisms in an alluvial mainstem river in northern Japan, where the tributary disappeared over the course of urban landscape transformation.

A 2.8-km long lowland segment of the mainstem gravel-bed river was examined for water properties and the river-riparian food web. In addition, watershed-wide water sampling was conducted to isotopically distinguish several types of groundwater that contributed to the hyporheic water in the study segment. There was a clear effect of altitude on the hydrogen/oxygen stable isotope ratios in the river water collected across the watershed.

Groundwater unique both in chemical sand isotopic signatures in several spots occurred within the study segment, and its properties resembled to and its upwelling locations matched groundwater from a tributary river whose surface channel has disappeared 60 years ago. Positive numerical increases in abundance and/or a sign of nitrogen transfer in river riparian communities (algae, invertebrates, and riparian trees) originating from groundwater high in nitrate with elevated nitrogen stable isotope ratios were found.

This study demonstrated that tributary groundwater with unique chemical properties manifested by an urban watershed river network continued to have cascading effects on biota across the river-riparian boundary in the mainstem river, even after urbanization transformed the tributary into a historically lost phantom river. We highlighted the legacy effects of landscape transformation in the subsurface domain and the significance of scrutinizing the past landscape and hydrological connectivity at the watershed scale in urban environments.

## Introduction

Urbanization is the major form of land transformation worldwide and is often associated with high environmental stresses on natural ecosystems and their services through climate change and pollution (Grimm et al., 2008). More than half of the world’s population lives in urban areas, and the trend of urban population growth continue, with an estimated 68% of the global population (9.7 billion in 2050) to be concentrated in urban areas by 2050 (United Nations, 2019, 2022). Therefore, for sustainable urban growth, understanding the key ecosystem structure and functions is imperative, building on which integrated policies to improve the lives of urban populations can be implemented. In rivers, urbanization results in changes in hydrology and the physical structure of stream channels and riparian zones, along with nutrient dynamics and biological communities (Pickett et al., 2011). However, studies on urban rivers are largely limited to reach-scale monitoring, requiring the inclusion of ecosystem perspectives at larger segment and watershed scales (Pickett et al., 2011). Urban landscapes are transformed by replacing natural river channels with completely new waterways, piping old channels, or diverting a part of water to elsewhere (Grimm et al., 2008), but subsurface hydrological connectivity is not necessarily equally modified. How and whether the subsurface domains of the original waterways and aquifers connecting them (a phantom of historical landscape) are functional is not known.

Longitudinal (upstream–downstream), lateral (river–riparian), and vertical (river–riverbed) dimensions characterize riverine ecosystems (Stanford and Ward, 1993). Vertically, the zones of surface water and groundwater mixing (hyporheic zones) play important roles in nutrient transformation, habitat provisioning, and temperature attenuation (Boulton et al., 1998; Boulton, 2007; Boulton et al., 2010). The hydrological exchange between surface water and groundwater is a key determinant of the ecological characteristics of the hyporheic zone, the magnitude and direction of which are strongly determined by the vertical hydraulic gradient and the substratum permeability (Brunke & Gonser, 1997; Gooseff, 2010). Consequently, the characteristics of hyporheic water are spatially diverse, and their physicochemical properties are affected by its sources, pathways, travel distance, and groundwater characteristics at multiple spatial scales. At a relatively small spatial scale, surface water tends to percolate into the riverbed substratum and upwells in immediate downstream areas within each geomorphic unit (Valett et al., 1994; Tonina & Buffington, 2007). River water may permeate at the upstream segment, travelling for several kilometers as subsurface flow in aquifer conduits and upwell at the downstream end of the alluvial fan or valley (Brunke & Gonser, 1997; Marmonier et al., 2020). Groundwater originate not only from infiltrated surface water that recharges aquifers but also from sources beyond focal river channels, such as other rivers and tributary catchments, and aquifers connecting them (Marmonier et al., 2020), underscoring the spatially explicit understanding of hydrological processes connecting groundwater and surface water at the watershed scale.

The ecological importance of the hyporheic zone can extend to surface water habitats and organisms in rivers. The contents and state of nutrients and organic matter in hyporheic flow are spatio-temporally variable, partially dependent on groundwater properties, and possess distinguished physical and chemical signatures relative to surface waters (Findlay, 1995; Brunke & Gonser, 1997). As a result, upwelling water can provide elements and environments that influence aquatic organisms, such as dissolved nitrogen and temperature heterogeneity. Dissolved nutrients are among the most important factors in the primary production in rivers (Biggs & Close, 1989). Patches with high nutrient supplies can be formed in upwelling zones, which enrich benthic algae biomass and accelerate recovery from disturbance (Valett et al., 1994; Wyatt et al., 2008). Through dampened heat exchange with the atmosphere in contact with groundwater and thermally stable environments, upwelling zones also provide high-quality spawning beds for salmon (Aruga et al., 2023). In addition, because the growth rate and/or abundance of aquatic insect larvae are affected by accumulated temperature, quality, and quantity of food resources (Sweeney & Vannote, 1982; Quinn et al., 1997), upwelling zones with thermally distinct habitats and/or abundant primary production may provide habitats for higher diversity and density of stream invertebrates (Marmonier et al., 2020). These increasingly known connections between subsurface processes and river ecosystems have rarely been investigated in the watershed landscape context, and none have been investigated in urban settings.

Live preys for consumers or the consumers themselves move across ecosystem boundaries and have reciprocal effects on each other as spatial resource subsidies (Polis et al., 1997). Streams and riparian zones are model systems in which two adjoining systems are trophically coupled, which play an important role in maintaining ecological stability and biological diversity (Baxter et al., 2005). In temperate headwater rivers, insects that fall from land to rivers provide substantial energy budgets for predatory fish (Kawaguchi & Nakano, 2001), and aquatic insects emerging from rivers to land provide food resources for riparian consumers (Nakano & Murakami, 2001; Fukui et al., 2006). In gravel-bed rivers, riparian arthropods, primarily spiders and ground beetles, dominate the riparian community in gravel-bar rivers (Framenau et al., 2002) and preferentially feed on emerging aquatic insects (Paetzold et al., 2005). In lowland rivers, furthermore, anthropogenic changes in river productivity and resource subsidies are common and nutrient enrichment of surface water and heavy metal pollution affect riparian consumers (Walters et al., 2008; Terui et al., 2018). Uno and Power (2015) demonstrated that vital spatial subsidies in river riparian interactions also occur via mainstem-tributary linkages in a river network. However, resource connectivity between land and groundwater via insect emergence is scarcely known (DelVecchia et al., 2016; Rahman et al., 2021), and the cascading effects of groundwater on riparian ecosystems from a watershed perspective are known only for riparian plants (Kuglerová et al., 2014).

This study examined the effects of tributary groundwater (GW) inflow on the response of riverine-riparian organisms in an alluvial mainstem river, where the tributary has disappeared over the course of urban landscape transformation. Previous investigations have reported that hyporheic water possesses distinct thermal characteristics and relatively high electrical conductivity (EC) in the Toyohira River, Hokkaido, Japan (Aruga et al., 2023; JNN, unpublished data), although its origin and formation mechanisms remain unknown. We hypothesized that this characteristic hyporheic water originated from groundwater contributed by the lost tributary river, and its effects would be reflected in biota on the riverbed and riparian zones of the mainstem river. We extensively utilized stable isotope ratio (SIR) measurements to elucidate the approximate source areas of GW and to trace nitrogen transfers from tributary GW to biota in the mainstem river. This was aided by the use of hydrogen/oxygen SIR, which can infer precipitation source areas based on altitude-dependent isotopic fractionation (McGill et al., 2020), and nitrogen SIR, whose variations together with carbon SIRs can indicate anthropogenic nitrogen pollution as well as riverine food web structure (Kaushal et al., 2011; Negishi et al., 2019; Alam et al., 2020). Because tributary catchments are located at relatively low elevations compared to the catchment of the mainstem river, we predicted that the source area of tributary-related GW could be distinguished from other sources. Furthermore, because benthic algae and aquatic insects that inhabit upwelling zones assimilate upwelling hyporheic water, we predicted that riverine organisms would have different SIR in the upwelling zone. Since elevated nutrients enrich primary production (Wyatt et al., 2008), we predicted that riparian invertebrates would also have a higher abundance and dependence on river-derived resources in the upwelling zone because of the higher abundance of emerging aquatic insects as prey (Terui et al., 2018).

## Methods

### Study site

The field study was conducted between 2012 and 2017 in the watershed of Toyohira River, which originate at Sapporo dake mountain (1440 m above sea level) flows through Sapporo City and is a tributary of the Ishikari River with 72.5 km long and a drainage area of 902 km^2^ (Fig. 1). Most of the Toyohira River watershed (824 km^2^, 91%) is located in Sapporo City, and 73.5% of Sapporo City belongs to the Toyohira River watershed (SM1). The segment is located in the lower part of the Toyohira Alluvial Fan, where alluvial deposits extend vertically up to several hundred metres (Sakata & Ikeda, 2013). Six major sub-basins comprise the Toyohira River watershed: the Toyohira River main-stem basin, Makomanai River basin, Shojin River basin, Motsukisamu River basin, Tsukisamu River basin, and Atsubetsu River basin. A 2.8-km long lowland study segment of the Toyohira River (approximately 10 m above sea level) was intensively examined because the presence of distinct groundwater was reported (Aruga et al., 2023). This segment currently provides >95% of Chum salmon (*Oncorhynchus keta*) spawning sites in the Toyohira River (Aruga et al., 2023). Among the major subbasins in the contemporary landscape, the Toyohira River mainstem, Makokanai River, and Shojin River basins contributed to the study segment via respective river channels and the Toyohira River channel.

**Figure 1.**
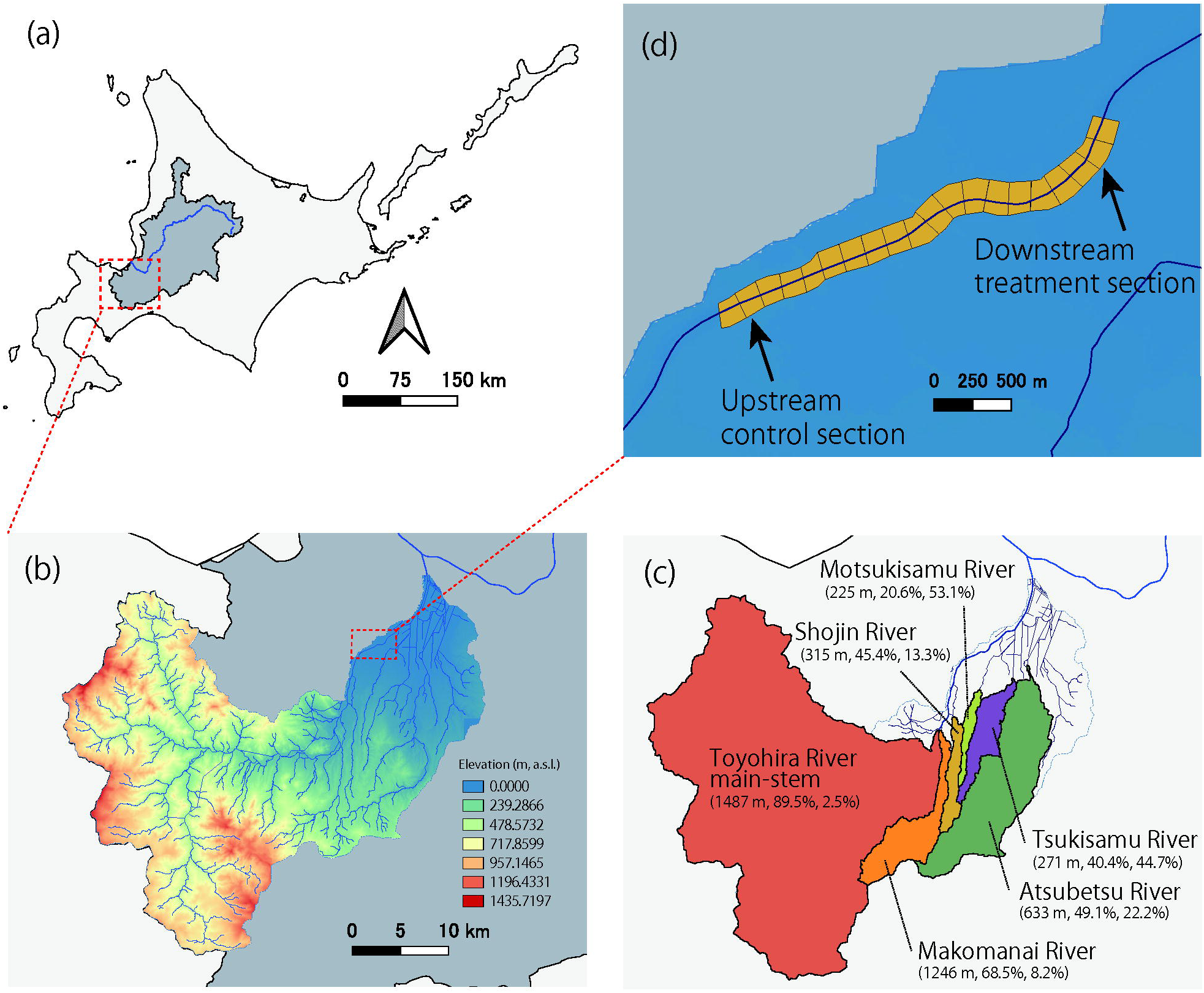
Map showing the study area in the Ishikari River watershed system (a), Toyohira River watershed (b), major sub-basin systems in the Toyohira River watershed (c), and the study segment with intensively monitored sections (d). In (b), color gradation indicates elevation (m above sea level), with a channel network in blue lines. In (c), the maximum elevation, proportion of forested land, and proportion of human land use (agriculture and buildups) are shown in brackets. In (d), the yellow-colored polygon shows 150-m compartments of the river channel used to organize data (see the text).

Cotemporary land cover in the Toyohira River mainstem subbasin is dominated by forests (89.5%), whereas land cover in other smaller basins at lower elevations is dominated by agriculture and buildups (up to 53.1%) (Fig. 1). Historically, landscape features of lowland areas of the watershed have transformed from agriculture-dominated ones to buildups-dominated ones with the population increases over the last 60 years; population of the city has grown by 800% to the current estimates of approximately 1.97 million in 2023 (SM2–4). Through urban growth, a tributary river that was present 60 years ago and appeared to have flowed into the area of the study segment disappeared (SM2-5). Although the natural river landscape of the studied segment was typical of gravel-bed rivers with alternating gravel bars, installations of embankments and groundsills, together with channel straightening over the past 100 years, transformed the river into a narrower one with limited lateral channel migration (SM5; Aruga et al., 2023). Mean annual flow rates and mean annual maximum flow rate (Kariki gauging station of MLIT, ca.100 m downstream of the study segment) over the past several decades were 26 and 440 m^3^/s, respectively.

### Detection of linkage between groundwater and benthic food web

Along the 2.8-km study segment, the relationship between hyporheic water characteristics and the benthic food web was assessed by examining the nitrogen SIR of benthic organisms. We calculated the deviations of EC in 20- to 30-cm deep hyporheic water relative to surface water based on existing data obtained in 2016 at 50-m intervals throughout the study segment as a proxy for targeted upwelling GW occurrence (SM6). From July 23 to August 3, 2015, herbivorous aquatic insect larvae (Heptageniidate spp.) and periphytic algae were collected at locations that approximately corresponded to the EC measurement locations. At each location, at least three cobbles were brushed to collect pooled samples of periphyton from the upper side of cobbles, and three to six individuals of Heptageniidate spp. were collected (182 individuals in total). The samples were dried in an oven at 60°C for 48 h, ground into a fine powder, and then packed into tin capsules. Carbon/nitrogen (C/N) SIR analysis was conducted using a mass spectrometer (MAT 252, Thermo Electron, Bremen, GER) as previously described (Alam et al., 2020). The carbon/nitrogen SIRs were ^13^C/^12^C and ^15^N/^14^N, expressed as δ^13^C and δ^15^N, respectively.

### Water characterizations within the river channel

Two 250-m long intensive study reaches were set up based on the levels of targeted upwelling GW occurrence: a treatment reach with a high level of GW occurrence and a control reach with low such influences. Five sections were set up every 50 m in each reach, with three sampling points 15-m distance from each other in each section. The two reaches were similar to each other in depth (mean ±SDs), and the flow velocity with the treatment reach was slightly deeper (44.51 cm±9.07 cm) than the control reaches (32.76 cm±10.22 cm) (SM7).

On February 1–6 and August 9, 2016, water samples were collected from both the surface and the hyporheic zones at each sampling point. We used the piezometer method (as mentioned above) to collect hyporheic water from a depth of approximately 20 cm in the riverbed and measured EC, pH, water temperature, and dissolved oxygen (DO) using portable probes (WM-32EP, DKK-KOA Co., TX 10, Yokogawa Meter & Instruments Co., and HQ 40 d, HACH Co.). Additionally, we measured these parameters for water seepage from the riverbank along the treatment reach five times (August 2012 and October 2014, and three times in October 2016). Water samples were preserved in airtight glass vials and polyethylene bottles, brought back to the laboratory, and preserved in a cool dark place for later analyses of SIRs of H_2_O as well as chemical constituents.

SIRs of hyporheic water and surface water were measured using gas bench (Gas Bench II, Thermo Fisher Scientific, Bremen, GER) and mass spectrometer (MAT 253, Thermo Electron, Bremen, GER). For water stable isotope analysis, the equilibrium method was used according to the following equation: δ= (R_sample_ / R_SMOW_ −1) × 1000 (‰). Here, δ indicates δ^18^O or δD and R indicates the isotopic ratio ^18^O/^16^O or D/H (^2^H/H). The measurement accuracies were ±0.05‰ for δ^18^O and ±0.05‰ for δD. The secondary standard SP2 (δ^18^O: −12.42‰, δD: −78.4‰) and DS2 (δ^18^O: −29.5‰, δD: −224.4‰), manufactured by the International Atomic Energy Agency (IAEA) and tested by VSMOW (Vienna SMOW) were used to correct the results. The major ions (NO_3_^−^, SO_4_^2−^, and Cl^−^) of water was measured using an ion analyzer (IA-300, DKK-KOA Co.).

### Water and land use characterizations at the watershed scale

Surface water was collected at 75 locations (115 samples) at different elevations, which is representative of potential sources of water to downstream study sections in different seasons between August 2012 and October 2016 (Table 1 and SM8). Base-flow river water samples were collected in airtight glass vials for later SIR analyses. Contributing catchment for each sampling location was delineated and their average and outlet elevation above the sea were obtained using 5- to 10-m resolution DEM data (Geospatial Information Authority of Japan) and GIS (QGIS, version 3.16.12). Oxygen/hydrogen SIR measurements were performed as described above.

**Table 1.**
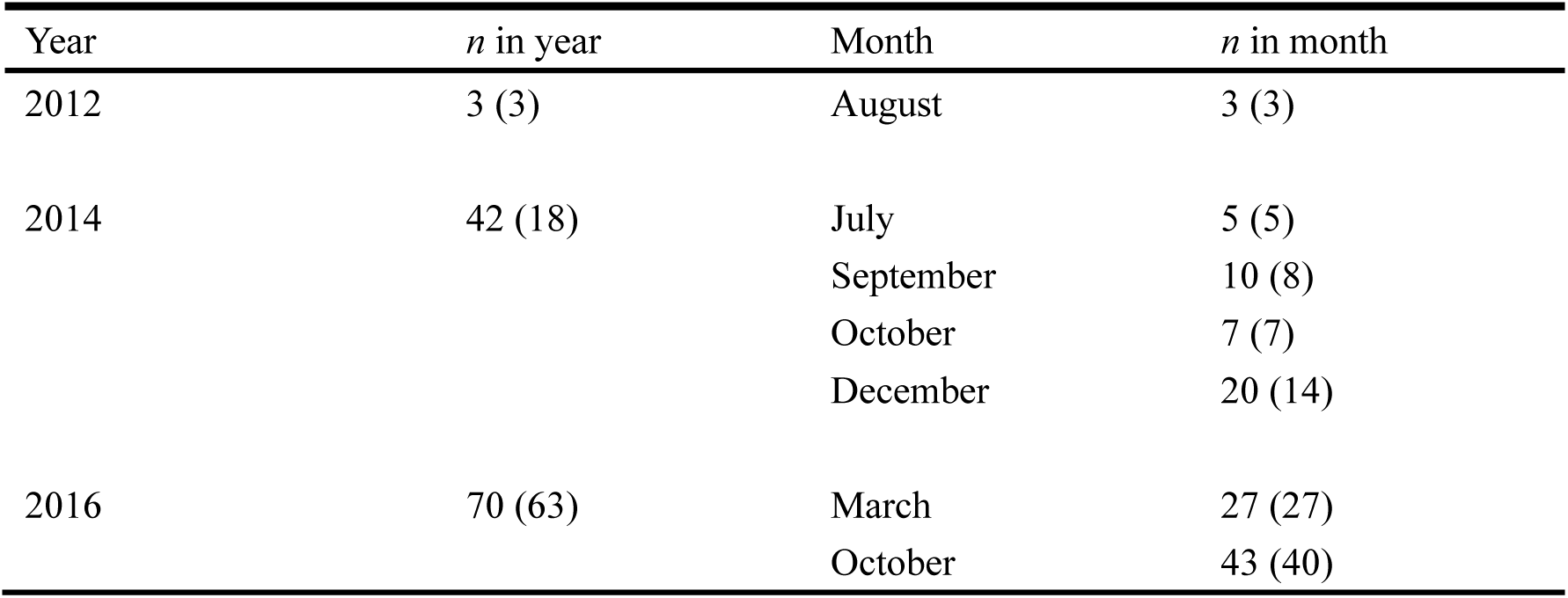
Sample size (*n*) of water collected for oxygen/hydrogen stable isotope ratio analyses at different locations of Toyohira River watershed, for each year and each month in each year. Numbers in brackets denote the number of sites visited, which was less than total sample size in some cases because some sites were sampled multiple times in the same month and year.

| Year | $n$ in year | Month | $n$ in month |
| --- | --- | --- | --- |
| 2012 | 3 (3) | August | 3 (3) |
| 2014 | 42 (18) | July | 5 (5) |
|  |  | September | 10 (8) |
|  |  | October | 7 (7) |
|  |  | December | 20 (14) |
| 2016 | 70 (63) | March | 27 (27) |
|  |  | October | 43 (40) |

### Benthic and riparian community

Abundances of benthic periphyton, invertebrates, and riparian arthropods were examined in the two reaches across different seasons. Two types of periphyton samples were collected by scrubbing three cobbles with a brush within a 100 cm^2^ area of each cobble (February 2, July 21, and Oct 19, 2016) and by setting up three unglazed ceramic tiles (100 cm^2^ each) (21 July, 2016) at each sampling point, followed by scrubbing as done for cobbles three weeks later. Benthic invertebrates were collected twice (February 2 and August 8, 2016) using a D-frame net (frame height 33 cm × bottom width 38 cm × net depth 45 cm, 350 μm mesh size) with one personnel disturbing the substratum immediately upstream (35 cm ×35 cm) at each point for 1 min. Riparian invertebrates were caught by burying plastic cups (diameter, 8 cm; depth, 15 cm) flush with the ground 1–2 m away perpendicularly from the river shore and longitudinally 10 m away from each other for 48-96 hours in the gravel bar. Approximately 50 ml of 20% propylene glycol was added to each cup during sampling. Pitfall sampling was repeated twice (July 23–25, 2016, August 5–9, 2017). Benthic invertebrate samples were immediately fixed with 75% ethanol, whereas riparian invertebrates were frozen.

The ash-free dry mass (AFDM) and chlorophyll *a* content of periphyton were measured as proxies for algal biomass. Periphyton samples were dried in an oven at 60°C for 48 h, weighed, combusted at 550°C for 3 h, and re-weighed to calculate AFDM per unit area (mg/m^2^). To measure algal chlorophyll *a*, extraction was conducted by filtering the periphyton samples using a glass fiber filter (GS-25, ADVANTEC Co.), dipping the filter in 90% acetone, and leaving it for 24 h in a cold dark place. Chlorophyll a concentration was measured by the SCOR-UNESCO (1966) method using an ultraviolet-visible spectrophotometer (UVmini-1240, SHIMADZU Co.) and converted to biomass per unit area (mg/m^2^).

Benthic invertebrates were identified mostly to the family level according to Maruyama and Takai (2000) and Kawai and Tanida (2018) using microscopy, and the number of each taxon was recorded. Riparian invertebrates were also identified to numerically dominant major groups (Carabidae, Araniae, Formicidae, and others) and enumerated.

For the food web analyses, benthic periphyton and insects were collected separately on July 23–28, November 3, 2015, and February 1–6 and July 21, 2016. Periphyton samples were scrubbed off from three cobbles with a brush randomly selected within each 50-m sub-sections (n = 20 for control reach and n = 20 for treatment reach), whereas three individuals of herbivorous Heptageniidae spp. were collected in each subsection (n = 60 for control and treatment reaches). Willow leaves from riparian forests were collected on October 19, 2016. In each 50-m section of the two reaches, four to five willow spp. trees were randomly chosen, and three to five leaves from each were collected as individual samples (n = 20 for control reach and n = 25 for treatment reach). Numerically dominant *Lithochlaenius noguchii* in riparian arthropods were collected from the above-mentioned individuals collected in pitfall traps after their abundance was recorded (n = 29 for control reach and n = 28 for treatment reach). C/N SIRs were measured as described above.

### Statistical analyses

The relationship between EC deviation and the nitrogen SIR of organisms was examined by developing generalized linear mixed models (GLMMs). Nitrogen SIR was the response variable, with EC deviation as the explanatory variable, and sample type as the random factor. The measured values at each 50-m sampling point were averaged for each 150-m distance because the two data could be spatially paired accurately at this spatial scale (Fig. 1d).

The water quality characteristics of four water types (surface and hyporheic water in two reaches) were characterized by principal component analysis (PCA) using a total of nine parameters for the two seasons.

Elevation effects on oxygen/hydrogen SRI were examined by developing GLMMs using δO as a response variable and catchment-average elevation as an explanatory variable, with different sampling occasions as a random factor. δ^18^O was used in the modelling because there was a strong correlation between δ^18^O and δD (Pearson’s correlation, p<0.001, r=0.95, n=120). Based on the GLMMs between elevation and δD, the elevation of the potential source area of water collected intensively in the monitored reaches and seepage water was estimated. The estimated elevation was then compared among the five water types developing generalized linear models (GLM) with elevation as a response variable and water type as an explanatory variable. When there was a statistically significant effect of the water type, multiple comparisons were performed to determine group differences.

Differences in algal biomass and invertebrate abundance among the reaches were examined by developing GLMMs using reach type as an explanatory factor. Both season and method type were included as random factors for algal biomass. For invertebrates, taxonomic group was also included as a random factor.

The C/N SIRs of aquatic and riparian organisms were compared between reach types using permutation multivariate analyses of variance (PERMANOVA) with taxonomic groups as a random factor. Furthermore, the contributions of aquatic resources to terrestrial consumers (*L. noguchii*) were estimated using Bayesian isotope mixing analyses separately for the two reach types, as done in previous studies (Alam et al., 2020). Heptageniidae spp. were used as an aquatic resource, whereas primary and secondary consumers relying on willow spp. as a basal resource were used as terrestrial resources. Primary and secondary consumers’ isotope signatures in riparian community were estimated by applying one and two trophic level equivalent mean isotope fractionations: 0.1‰ (C) and 3.4‰ (N) for one and 0.2‰ (C) and 6.8‰ (N) for two levels. Because the proportion of primary and secondary consumers of terrestrial origin consumed proportionately by carabid beetles was unknown, we estimated the contribution for three scenarios:100% primary consumers, 100% secondary consumers, and 50 to 50% of each, in which the fractionation factor was proportionately adjusted according to assumed mixture ratios.

The R software (R Core Team, 2022) was used to perform all statistical analyses. The significance level was set at p=0.05. Multiple comparisons were also performed. In the adonis package for PERMANOVA, Euclidian distance was used to distinguish groups.

## Results

There was a significant positive relationship between the differences in EC between the two zones and the nitrogen SIR (Table 2; Fig. 2). Two reaches were selected based on this relationship, with the treatment reach being higher in EC difference and nitrogen SIR compared to those in the control reach.

**Figure 2.**
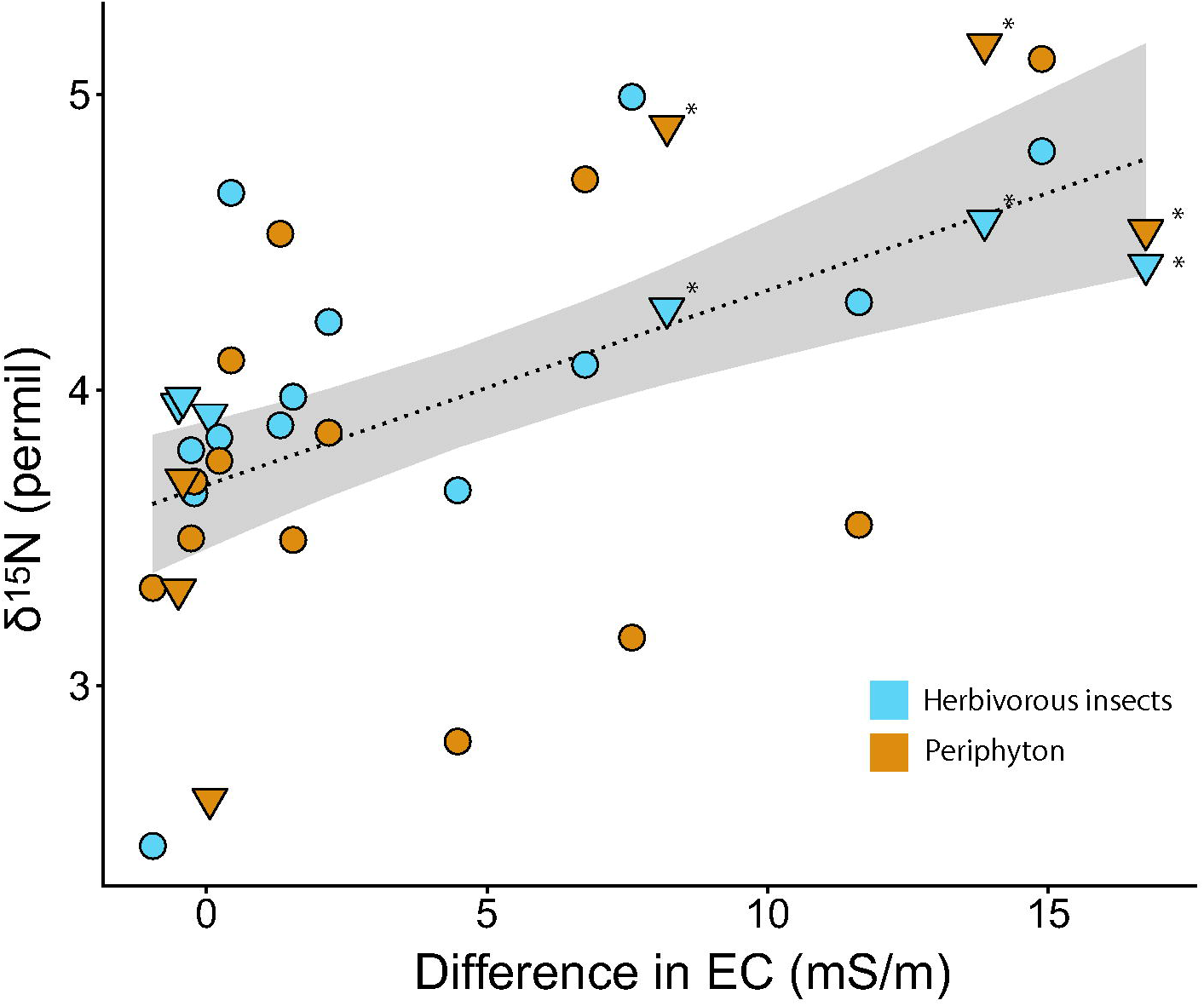
Relationship between the upwelling groundwater index (differences in electrical conductivity of hyporheic water relative to that of the surface zone) and nitrogen stable isotope ratios of herbivorous insects (Heptageniidae spp.) and periphyton. Each data point represents the mean for each 150-m study comportments in the study segment. Dotted lines and grey clouds denote the regression model and its 95% confidence intervals, respectively. Triangle symbols indicate samples collected from two reaches selected for later intensive monitoring; those with asterisks are from the reach assigned as treatment reach.

**Table 2.**
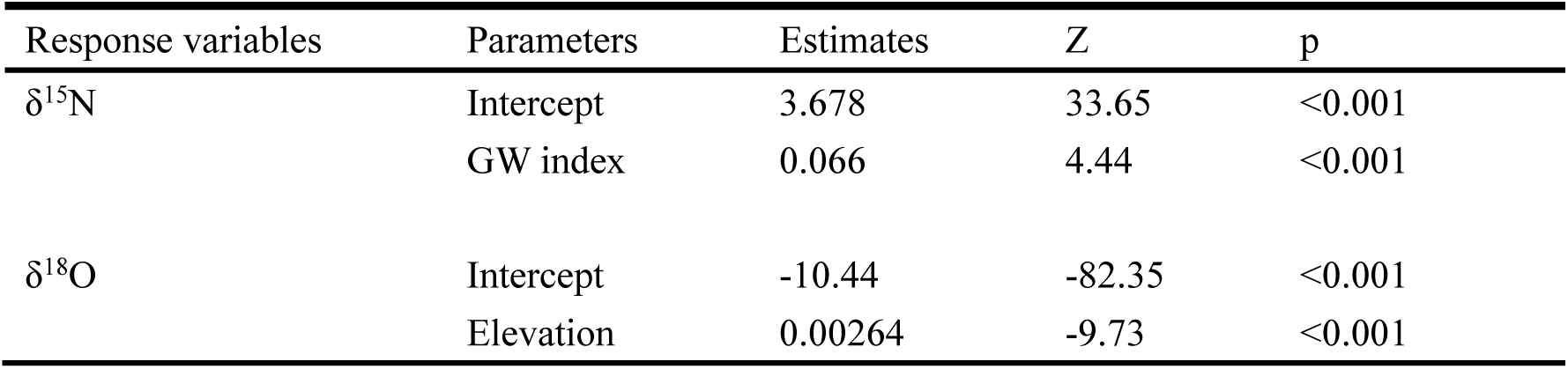
Results of generalized mixed linear models testing effects of the targeted upwelling groundwater index (GW index) (differences in electrical conductivity of water between hyporheic zone and surface water zone) on nitrogen stable isotope ratios (δ^15^N) of herbivorous insects (Heptageniidae spp.) and periphyton, and the effects of catchment elevation on the oxygen stable isotope ratios (δ^18^O) of water.

The water quality characteristics in the study reaches were summarized into two principal component scores, with approximately 75% of the variation explained, of which the first score accounted for >58% of the variation (Table 3). On both occasions, the first score was highly and positively correlated with EC (and other chemical constituents such as NO_3_^−^, SO_4_^2−^, and Cl^−^) and oxygen/hydrogen SIRs. A strong negative association was observed between the DO and pH. In two-dimensional spaces based on the first two scores, water quality was different only in hyporheic water in the treatment reach on both sampling occasions, particularly along the PC1 axis (Fig. 3). Seepage water was characterized with water quality properties and hydrogen/oxygen SIR most similar to hyporheic water in the treatment reach (Table 4).

**Figure 3.**
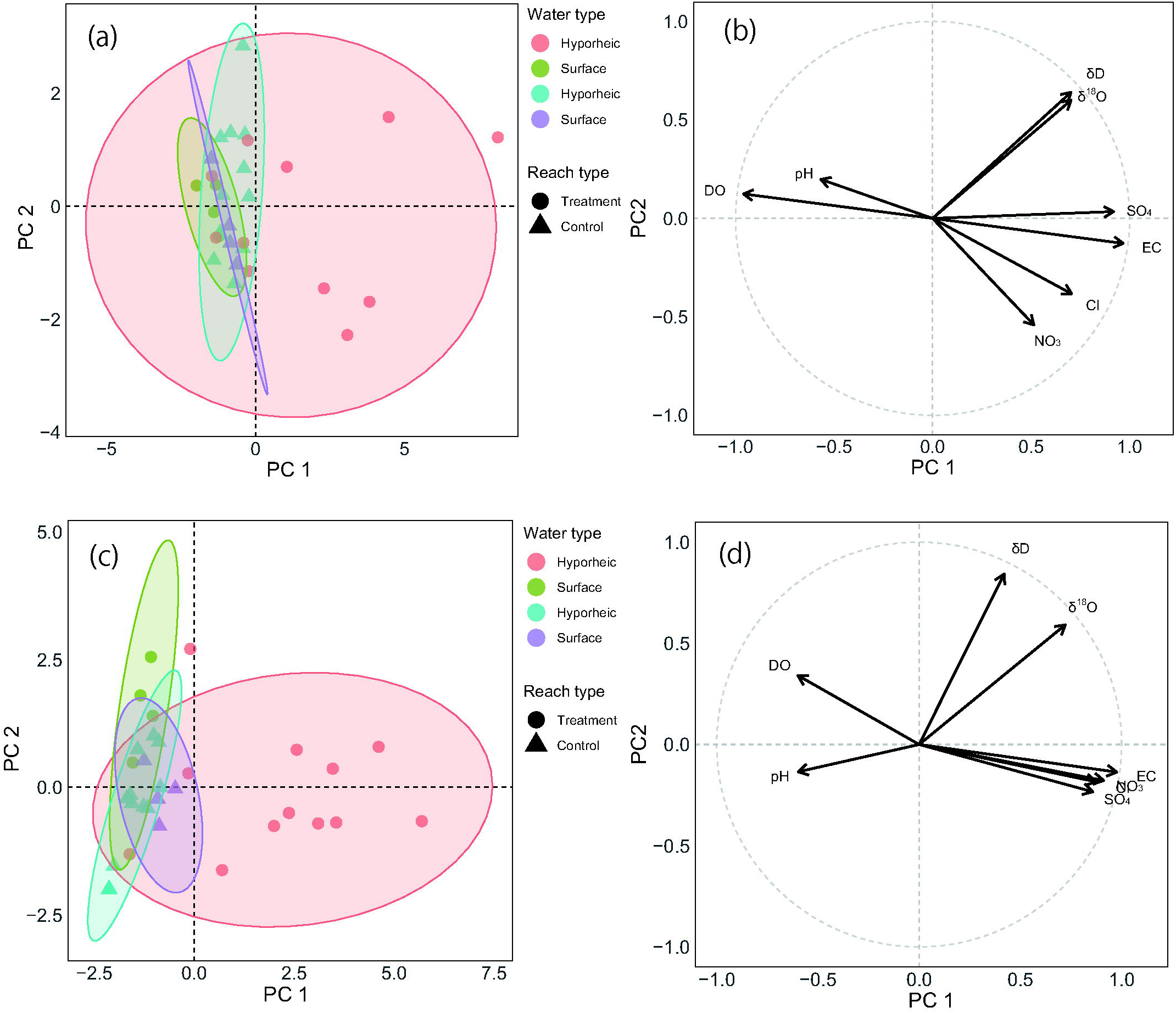
Principal component (PC) scores for four types of water in February 2016 (a) and August 2016 (b). Loadings of each environmental variable to the PC scores are shown for February 2016 (c) and August 2016 (d).

**Table 3.** Results of principal component analyses to characterize water in winter (February) and summer (August). Environmental factors were shown in relation to PC scores with their eigenvalues and variance explained (%). Two seasons were analyzed separately. Abbreviated environmental factors: electrical conductivity (EC); dissolved oxygen (DO); nitrate (NO_3_), chloride (Cl), sulfate (SO_4_), and stable isotope ratios of oxygen and hydrogen (δ^18^O and δD). Pearson’s correlation coefficient with respective PC scores were shown with their statistical significance: *<0.05; **<0.01; ***<0.001.

|  | Winter |  | Summer |  |
| --- | --- | --- | --- | --- |
| Principal components | PC1 | PC2 | PC1 | PC2 |
| Eigenvalues | 4.62 | 1.24 | 4.55 | 1.29 |
| Variance explained (%) | 59.64 | 16.02 | 58.68 | 16.66 |
| Environmental factor |  |  |  |  |
| EC | 0.97*** | 0.13 | 0.98*** | 0.14 |
| pH | -0.57*** | -0.20 | -0.60*** | 0.13 |
| DO | -0.96*** | -0.12 | -0.60*** | -0.34 |
| NO<sub>3</sub> | 0.51** | 0.54** | 0.91*** | 0.18 |
| Cl | 0.71*** | 0.38 | 0.87*** | 0.18 |
| SO<sub>4</sub> | 0.92*** | -0.03 | 0.86*** | 0.23 |
| δ<sup>18</sup>O | 0.70*** | -0.60*** | 0.72*** | -0.59*** |
| δD | 0.70*** | -0.64*** | 0.42* | -0.84*** |

**Table 4.**
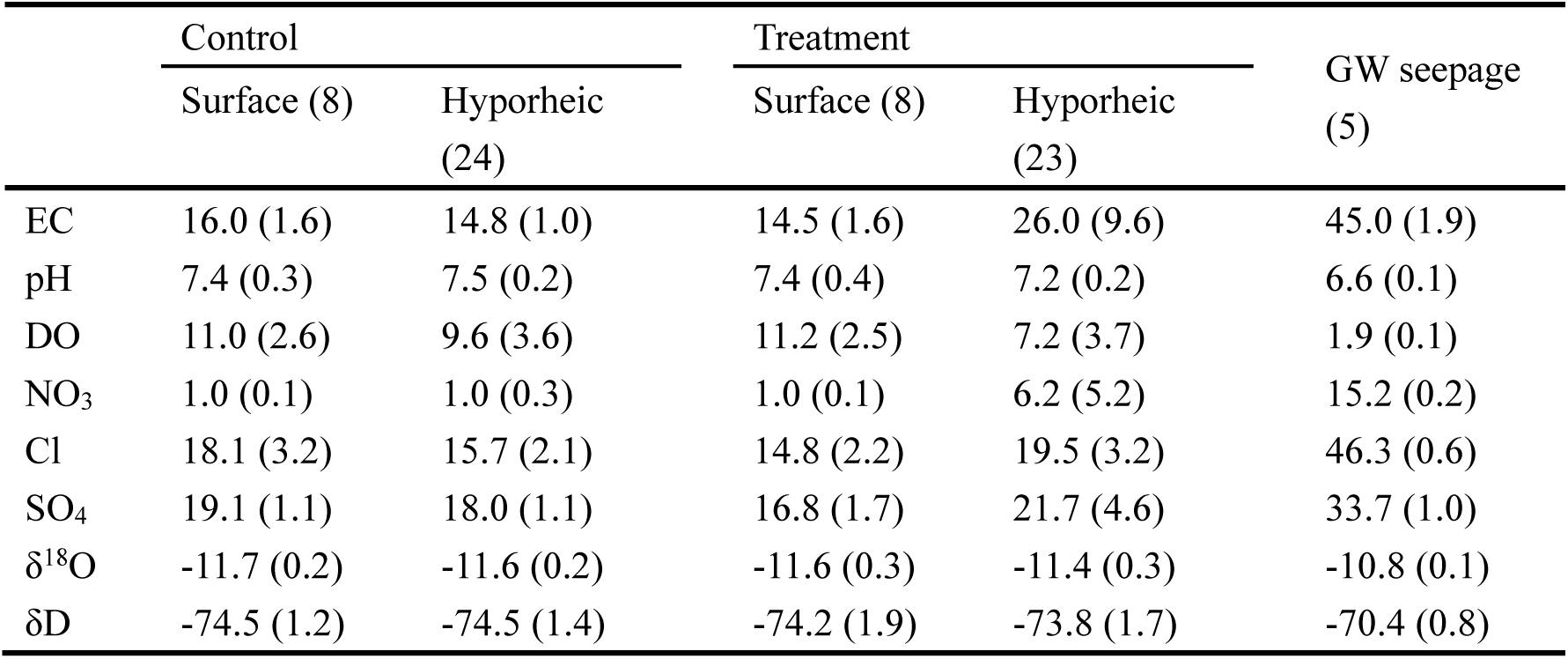
Means (±SDs) of physico-chemical characteristics and hydrogen/oxygen stable isotope ratios of water collected in the study reaches and groundwater seepage from the river bank. The numbers in brackets indicate sample sizes. Abbreviations and units were: electrical conductivity (EC, mS/m); dissolved oxygen (DO, mg/L); nitrate (NO_3_, mg/L), chloride (Cl, mg/L), sulfate (SO_4_, mg/L), and stable isotope ratios of oxygen and hydrogen (δ^18^O and δD, permil).

|  | Control |  | Treatment |  | GW seepage<br>(5) |
| --- | --- | --- | --- | --- | --- |
|  | Surface (8) | Hyporheic<br>(24) | Surface (8) | Hyporheic<br>(23) |  |
| EC | 16.0 (1.6) | 14.8 (1.0) | 14.5 (1.6) | 26.0 (9.6) | 45.0 (1.9) |
| pH | 7.4 (0.3) | 7.5 (0.2) | 7.4 (0.4) | 7.2 (0.2) | 6.6 (0.1) |
| DO | 11.0 (2.6) | 9.6 (3.6) | 11.2 (2.5) | 7.2 (3.7) | 1.9 (0.1) |
| $\text{NO}_3$ | 1.0 (0.1) | 1.0 (0.3) | 1.0 (0.1) | 6.2 (5.2) | 15.2 (0.2) |
| Cl | 18.1 (3.2) | 15.7 (2.1) | 14.8 (2.2) | 19.5 (3.2) | 46.3 (0.6) |
| $\text{SO}_4$ | 19.1 (1.1) | 18.0 (1.1) | 16.8 (1.7) | 21.7 (4.6) | 33.7 (1.0) |
| $\delta^{18}\text{O}$ | -11.7 (0.2) | -11.6 (0.2) | -11.6 (0.3) | -11.4 (0.3) | -10.8 (0.1) |
| $\delta\text{D}$ | -74.5 (1.2) | -74.5 (1.4) | -74.2 (1.9) | -73.8 (1.7) | -70.4 (0.8) |

Elevation was negatively correlated with the oxygen SIR (Table 2; Fig. 4a). When examined in relation to water originating from different sub-basins, water collected in the Toyohira River mainstem sub-basin exhibited lower oxygen SIR relative to water collected in other basins, even at an elevation as low as the intensively monitored reaches (Fig. 4b). The oxygen SIR of seepage water was higher than that of water collected in the Toyohira River main-stem system and only overlapped with those of water originating from lower elevation subbasins.

**Figure 4.**
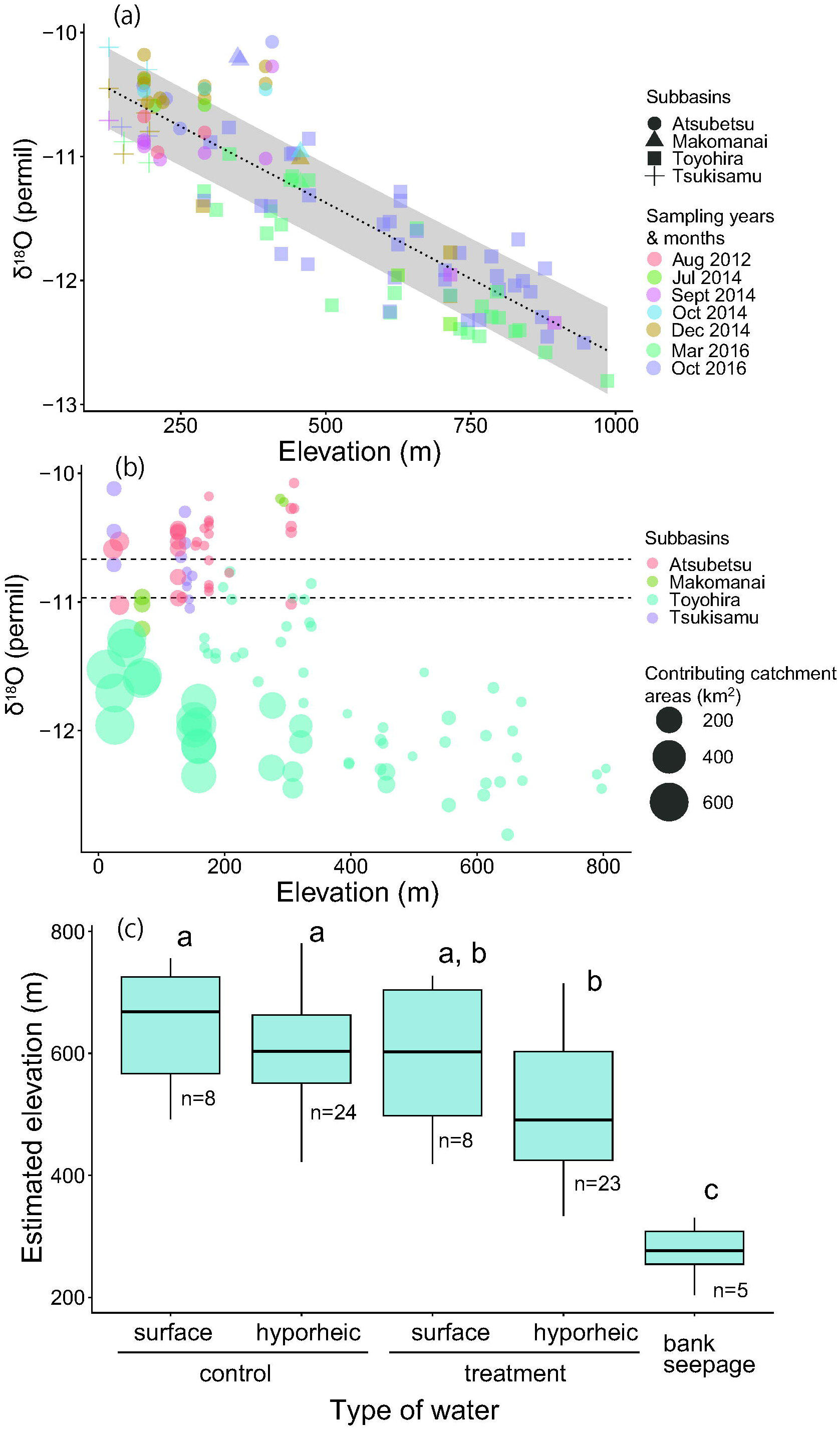
Relationship between the catchment average elevation of sampling sites and oxygen stable isotope ratio of water collected at different locations within the Toyohira River watershed (a). Biplot showing oxygen stable isotope ratios of water collected at different elevations in relation to their contributing catchment size and sub-basin category (b). Comparisons of estimated elevation of source area contributing to water collected in surface and hyporheic zones of two reaches and seepage water from the riverbank (c). In (a), the dotted line and gray clouds denote the linear regression line and 95% confidence intervals, respectively. In (b), the horizontal dotted lines denote the upper and lower ranges of the 95% confidence intervals of isotope ratios for water collected from seepage water from the riverbank. In (c), alphabetical letters denote multiple comparison results among groups; those accompanied by the same letter are statistically undistinguished.

The oxygen SIR significantly differed among the groups of different water types (GLM, p<0.001). Multiple comparison tests revealed that the water source elevations of three water types other than water from the hyporheic zone of the treatment reach were similar to each other (Fig. 4c). Furthermore, hyporheic water in the treatment reach was significantly different from that in the control reach at its source elevation. Overall, water from the hyporheic zone of the treatment reach was the lowest in terms of its source elevation, with a median elevation of approximately 510 m. Seepage water was estimated to have source elevation at 200–300 m.

A total of 9907 aquatic invertebrates belonging to 26 taxa were collected, and 1289 terrestrial invertebrates from nine taxa were sampled (Table 5). Among the carabid beetles, *L. noguchii* comprised 89.9% of the total abundance of this group. The abundance of all the numerically examined taxa differed between reach types (Table 6) and was higher in the treatment reach than in the control reach (Fig. 5). C/N SIRs of the organism groups differed between the reach types (p<0.001) (Fig. 6a). The contribution of aquatic resources to consumers was higher in the treatment reach than in the control reach under all three scenarios, especially in scenarios in which primary consumers provided 100% and 50% of the terrestrial resources to carabid beetles (Fig. 6b).

**Figure 5.**
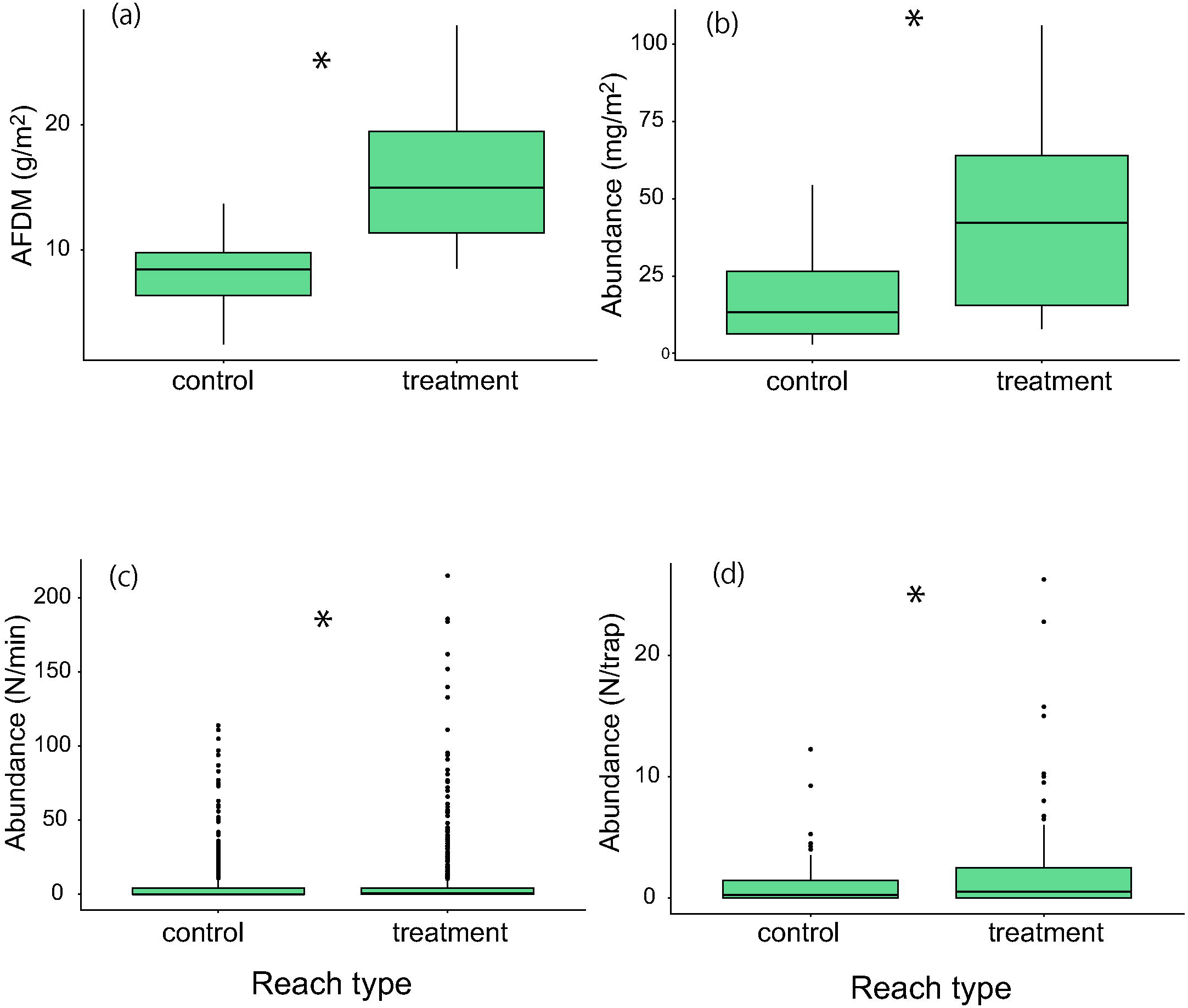
Comparison of ash-free dry mass (AFDM) (a) and chlorophyll *a* abundance (b) of periphyton as an index of algal biomass, and aquatic invertebrate abundance (c) and terrestrial arthropod abundance (d) between reach types. Asterisks denote statistically significant differences between the groups.

**Figure 6.**
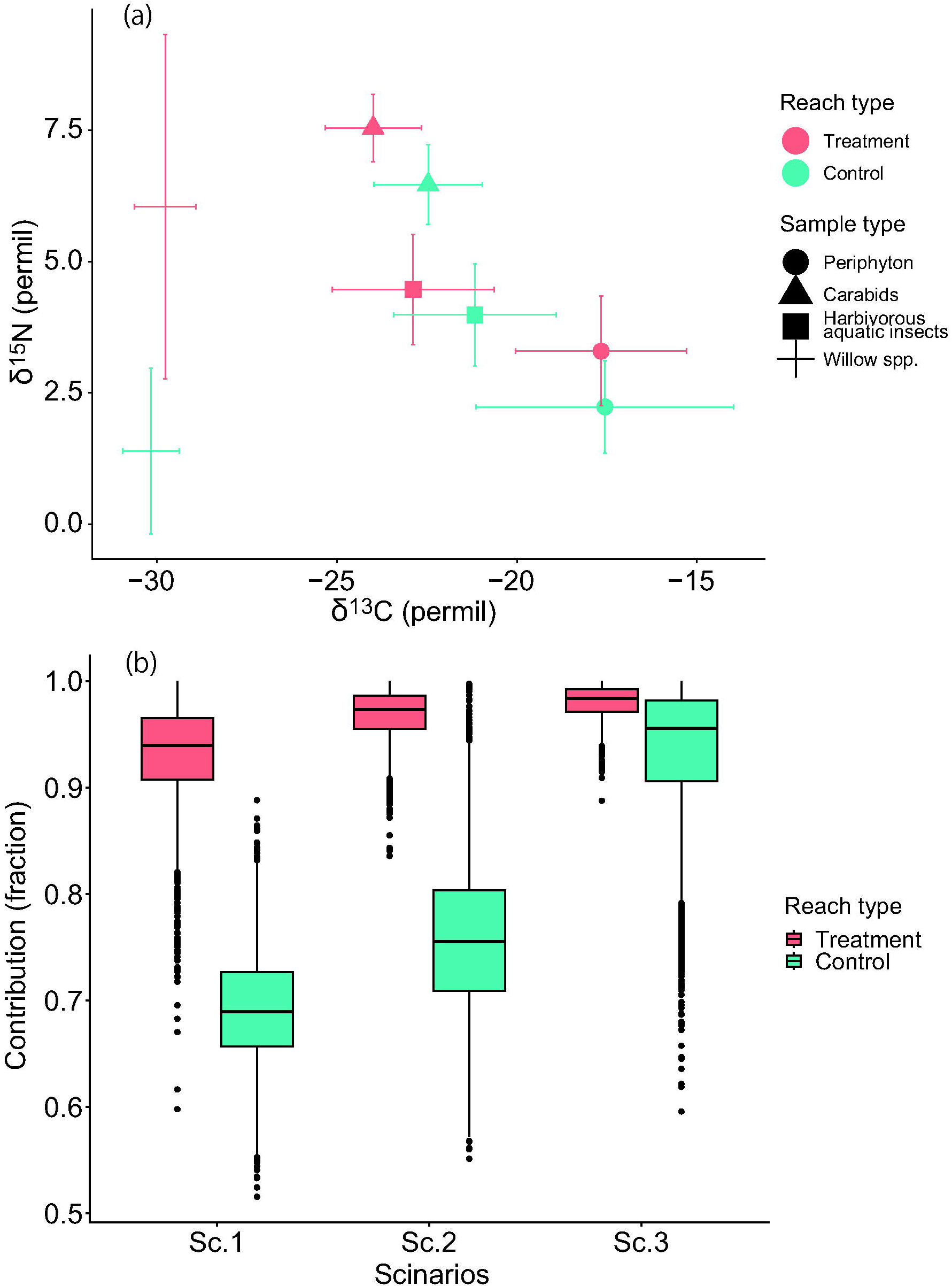
Carbon and nitrogen stable isotope ratios of four sample types in two reaches (a) and estimated contributions of aquatic resources to terrestrial consumers (Carabidae, *Lithochlaenius noguchii*) in the two reaches (b). In (a), the mean and standard deviation are plotted for each group. In (b), boxplots are based on data points (n=5000 for each group) generated using Bayesian isotope mixing models.

**Table 5.** Total abundance and mean (±SD) abundances of aquatic invertebrates (a) and terrestrial invertebrates (b) in the control and treatment reaches. Numbers in brackets of abundance denote the relative proportion of each taxon (%) whereas *n* indicates samples size.

| Taxa | Abundance | Control reach (n=27) | Treatment reach (n=26) |
| --- | --- | --- | --- |
| Chironomidae | 4311 (43.51) | 57.8 (32.1) | 87.9 (56.2) |
| Tipulidae | 1035 (10.45) | 13.4 (12.8) | 24.2 (25.1) |
| *Epeorus* | 1029 (10.39) | 20.0 (11.1) | 13.9 (12.7) |
| Hydropsyche | 841 (8.49) | 8.4 (12.4) | 17.0 (18.5) |
| Stenopsyche | 436 (4.40) | 5.0 (7.0) | 10.6 (10.9) |
| *Cincticostella* | 384 (3.88) | 5.3 (7.4) | 4.7 (9.5) |
| *Uracanthella* | 342 (3.45) | 4.8 (5.3) | 6.0 (7.4) |
| *Baetis* | 285 (2.88) | 7.0 (8.8) | 3.6 (4.3) |
| *Baetiella* | 232 (2.34) | 5.1 (9.3) | 3.6 (5.6) |
| *Paraleptophlebia* | 190 (1.92) | 3.3 (5.6) | 2.5 (3.1) |
| Perlodidae | 187 (1.89) | 2.8 (3.0) | 3.0 (2.1) |
| *Caenis* | 115 (1.16) | 1.9 (3.1) | 2.4 (2.4) |
| *Ephemerella* | 108 (1.09) | 1.6 (2.2) | 2.5 (4.1) |
| *Ecdyonurus* | 86 (0.87) | 2.0 (3.3) | 1.2 (2.0) |
| *Rhithorogena* | 59 (0.60) | 0.9 (1.8) | 0.8 (1.7) |
| *Ameletus* | 59 (0.60) | 0.6 (1.0) | 1.5 (2.9) |
| *Torleya* | 56 (0.57) | 1.0 (1.9) | 0.6 (1.1) |
| *Heptagenia* | 54 (0.55) | 0.6 (1.5) | 0.9 (1.7) |
| Elmidae | 25 (0.25) | 0.2 (0.6) | 0.8 (2.1) |
| *Drunella* | 21 (0.21) | 0.4 (0.7) | 0.2 (0.5) |
| Glossosomatidae | 15 (0.15) | 0.1 (0.4) | 0.4 (0.9) |
| Polycentropodidae | 14 (0.14) | 0.3 (0.8) | 0.3 (0.7) |
| Chloroperlidae | 9 (0.09) | 0.3 (0.6) | 0.1 (0.2) |
| *Rhyacophila* | 7 (0.07) | 0.1 (0.3) | 0.1 (0.3) |
| Amphipoda | 6 (0.06) | 0.0 (0.2) | 0.2 (0.6) |
| *Dolophilodes* | 1 (0.51) | 0.0 | 0.02 (0.1) |

| Taxa | Abundance | Control reach (n=22) | Treatment reach (n=22) |
| --- | --- | --- | --- |
| Formicidae | 448 (34.8) | 0.78 (0.99) | 4.59 (7.28) |
| Cardiophorinae | 321 (24.9) | 1.85 (2.73) | 1.99 (2.23) |
| *Lithochlaenius*<br>*noguchii* (Bates,<br>1873) | 260 (20.2) | 1.91 (1.70) | 2.09 (2.39) |
| Others | 68 (5.3) | 0.28 (0.65) | 0.53 (0.69) |
| Staphylinidae | 44 (3.4) | 0.03 (0.12) | 0.49 (0.78) |
| Carabidae spp. | 42 (3.3) | 0.15 (0.26) | 0.41 (0.49) |
| Araneae | 42 (3.3) | 0.28 (0.64) | 0.28 (0.44) |
| Dermaptera | 35 (2.7) | 0.00 | 0.40 (1.16) |
| *Brachinus scotomedes*<br>Redtenbacher | 29 (2.2) | 0.03 (0.12) | 0.50 (1.07) |

**Table 6.** Results of generalized mixed linear models testing effects of reach type on algal biomass, aquatic invertebrates, and terrestrial arthropods. For algal biomass, ash free dry mass (AFDM) and chlorophyl *a* (chl. *a*) abundance were used.

| Response variables | Parameters | Estimates | Z | p |
| --- | --- | --- | --- | --- |
| AFDM | Intercept | 0.8153 | 4.95 | <0.001 |
|  | Reach type | 0.7929 | 3.404 | <0.001 |
| chl. *a* | Intercept | 29.452 | 1.923 | 0.054 |
|  | Reach type | 24.59 | 3.793 | <0.001 |
| aquatic invertebrates | Intercept | 0.16401 | 0.359 | 0.72 |
|  | Reach type | 0.22985 | 11.136 | <0.001 |
| terrestrial arthropods | Intercept | -0.66448 | -0.996 | 0.32 |
|  | Reach type | 0.7905 | 13.129 | <0.001 |

## Discussion

Despite the increasingly recognized importance of tributary–mainstem connectivity as a node of spatial resource subsidy and hotspots of habitat heterogeneity, and the dendric watershed river network as a key in genetic diversity and population dynamics (Benda et al., 2004; Kelson et al., 2015; Uno & Power, 2015; Terui et al., 2018), the majority of urban stream ecological studies have examined reach-scale processes (Pickett et al., 2011). This study demonstrated the influence of subsurface connectivity of tributary flow to the mainstem river with a watershed perspective in an urban setting, filling the current knowledge gap. These findings largely support our initial predictions. First, hyporheic water in the study segment was unique in terms of water quality, and the SIR signature indicated high contributions of tributary inflow compared with surface water flowing from upstream. Second, the presence of this hyporheic water was associated with a higher abundance of riverine organisms, labelled with an elevated level of nitrogen SIR. Lastly, riparian invertebrates had a higher abundance and dependence on rivers, with high contributions of tributary-derived groundwater to their habitat.

The linear relationship between hyporheic water quality in the EC and nitrogen SIR in benthic communities indicated the presence and spatial variability of trophic linkages via nitrogen flux operating at the surface and subsurface interface. The nitrogen SIR of consumers increases in the presence of synthetic nitrogen (Kaushal et al. 2011; Negishi et al. 2019; Alam et al. 2020). The nitrogen SIR of organic matter and dissolved nitrogen are commonly elevated by anthropogenically polluted water. The mechanisms are complex, but the general cause is the addition or formation of nitrogen with an increased proportion of heavier SIR in particulate or dissolved matter by volatilization and denitrification (Botrel et al., 2014). Detailed water chemistry analyses in the two reaches clearly showed that the elevated EC in hyporheic water was partially driven by high levels of nitrate. The levels of nitrate observed were high and were comparable with the average levels reported in human-affected groundwater in farmlands and livestock fields in Japan (1.4 to 20 mg/L as NO_3−_N; 3.15 to 45 mg/L as nitrate) (Yabusaki, 2010). Globally, human activities, such as urbanization and agriculture, have elevated nitrate levels in water to comparable levels (Drake & Bauder, 2005; Negishi et al., 2019; Su et al., 2021). Thus, it is plausible that upwelling hyporheic water partially affected by human-introduced nitrogen was assimilated into the benthic community through primary producer uptake followed by consumer resource consumption, creating noticeable hotspots of trophic linkages in the lowland mainstem of the Toyohira River.

Critical knowledge of the hyporheic environment is the hydrological process that maintains groundwater component mixing with surface water because it helps to understand the spatial scale over which surface-to-subsurface connectivity operates, as well as the mechanisms behind the formation of physico-chemical properties of hyporheic water (Brunke & Gonser, 1997; Baxter & Hauer, 2000). In our case, the water SIR of hyporheic water provided keystone information regarding its origin and pathway, which was otherwise difficult to discern using only chemical constituents. For example, water chemistry has limited ability to reveal the origin of nitrate-rich hyporheic water. Nitrate-rich water can be generated in the riverbed when upwelling water undergoes mineralization of organic matter and/or nitrification of nitrogen compounds (Krause et al., 2013). Alternatively, nitrate-rich water with high nitrate loadings could be sourced from elsewhere (Negishi et al., 2019; Alam et al., 2020). Water SIR has been used as a powerful racer tool in appreciating hydrological cycles and pathways because atmospheric water vapor, precipitation, runoff, and groundwater can have temporally and spatially distinct values as a result of isotopic fractionations operating at local, regional, and continental scales (Gat, 1996). Hyporheic water in the treatment reach was not only rich in dissolved elements, but was also characterized by higher hydrogen SIR values compared to both types of water in the upstream control reach, which clearly indicated that the origin of groundwater was not infiltrated surface water into the river bed, which is similar to the case of Marmonier et al. (2020). We support the strength of an approach combining isotopic and water quality signatures for water source determination in the hydrological cycle (Torres-Martínez et al., 2020; Su et al., 2021).

Seepage water resembled hyporheic water in the treatment reach both in water chemistry and isotopic signatures, providing strong evidence that characteristic hyporheic water detected in the mainstem Toyohira River was a mixture of this seepage groundwater and surface water flowing down from upstream areas. A common application of water SIRs as a hydrological tool is the determination of source areas based on altitude-dependent water SIR signatures (McGill et al., 2020). Precipitation at high altitudes exhibit lower isotope ratios than that at low altitudes, which is known as the altitude (elevational) effect (Dansgaard, 1964; Clark & Fritz, 1997). In our system, this elevational effect was evident. The estimated source water altitude of more than 50% of the surface water samples collected in the mainstem of the Toyohira River was >600 m, reflecting the catchment’s altitude that extends >1400 m above sea level. In contrast, seepage water was estimated to have originated from a much lower altitude, which approximately matches the altitudes of smaller low-altitude sub-basins and rivers such as the Shojin, Motsukisamu, and Tsukisamu Rivers. Although there is currently no river channel that flows into the study segment, there is a channel and catchment whose channel merges with the mainstem at almost the exact location of the study segment. Thus, we believe that the surface catchment topography of this lost river still collects precipitation, and the resultant subsurface flow is connected to the mainstem. The low-lying areas of this lost catchment have abundant alluvial deposits from the Toyohira River (Sakata & Ikeda, 2013); thus, the subsurface zone is expected to be conducive to groundwater flow. High levels of nitrate with a sign of anthropogenic effects in seepage groundwater are reasonable because water goes through the areas with high population density and nitrogen sources, such as anthropogenically modified soil organic matter, sewer system leakages, and chemical fertilizers to crop fields.

Food web analyses in the two selected reaches revealed trophic cascading effects of groundwater extending to riparian consumers. CN SIR measurements showed that all four groups of organisms, including willows spp. in the treatment reach, assimilated tributary-derived groundwater, as seen in an elevated N SIR signature. Thus, dual-pathway trophic linkages between groundwater and riparian zones were demonstrated. This groundwater-derived nitrogen assimilation in aquatic consumers apparently co-occurred with the numerical response of aquatic consumers, probably because high nitrate levels acted as resource subsidies with bottom-up effects on primary producers and herbivorous insects (Negishi et al., 2019). Other functional feeding invertebrates presumably benefitted from resource subsidies because algae-rich periphyton layers were also inhabited by taxa such as the numerically dominant Chironomidae. Riparian plants use groundwater to maintain productivity and diversity (Dawson & Ehleringer, 1991; Jansson et al., 2007; Kuglerová et al., 2014). Our results were indicative of willow trees’ uptake of groundwater, although it remains unknown whether groundwater subsidy caused any changes in riparian tree productivity or whether there were cascading effects on terrestrial consumers propagating through terrestrial herbivores’ numerical responses. Riparian consumers increased not only their abundance but also their dependence on riverine prey, suggesting that the trophic cascading effects of tributary-derived groundwater might have been amplified disproportionately via a route of river to the riparian zone with an increased insect emergence (Terui et al., 2018).

In conclusion, this study demonstrated that tributary groundwater with unique chemical properties manifested by an urban watershed river network continued to have cascading effects on biota across the river-riparian boundary in the mainstem river, even after urbanization transformed the tributary into a historically lost phantom river. The legacy effects of past ecosystem structures on contemporary biological community structure and distribution are well known in freshwater systems (Harding et al., 1998; Negishi et al., 2014). Our findings extend these by highlighting the legacy effects of landscape transformation in the subsurface domain and the significance of scrutinizing the past landscape and hydrological connectivity at the watershed scale in urban environments. Furthermore, the discovery of functional subsurface connectivity in the river riparian food web provides insights into the habitats of other taxa. Specifically, this segment provides important spawning grounds for Salmonidae fish, and there are spots of high hyporheic water temperature that can affect the quality of the spawning habitat (Aruga et al., 2023). Although the current study did not report thermal heterogeneity in the mainstem river created by the presence of tributary groundwater, it is probable that some of the reported thermal heterogeneity in the area is attributable to the phantom tributary flow. Future studies, including fish habitat quality studies, warrant a more holistic understanding of the ecological functions of a lost river and critical knowledge for sustainably managing urban river networks.

## Supporting information

Supplementary materials all

## Acknowledgements

This study was partly supported by the research fund for the Tokachi and Ishikari rivers provided by MLIT (18056588) and JSPS KAKENHI (18H03408 and 18H03407).

## Ethical approval

No ethical violation was occurred in this research.

## Data availability statement

All the data used in the paper will be deposited in Dryad.

## Author contributions

JN Negishi: Conceptualization (equal), data collection (equal), data curation (equal), formal analysis, funding acquisition, and leading writing. YY Song, I Matsubara, and N Morisaki: Conceptualization (equal), data collection (equal), data curation (equal), preliminary analyses (equal). All the authors contributed critically to the drafts and approved the final manuscript for publication.

