## Supplementary materials all for "An urbanized phantom tributary subsidizes river-riparian communities of mainstem gravel-bed river"

Article tile: Research paper

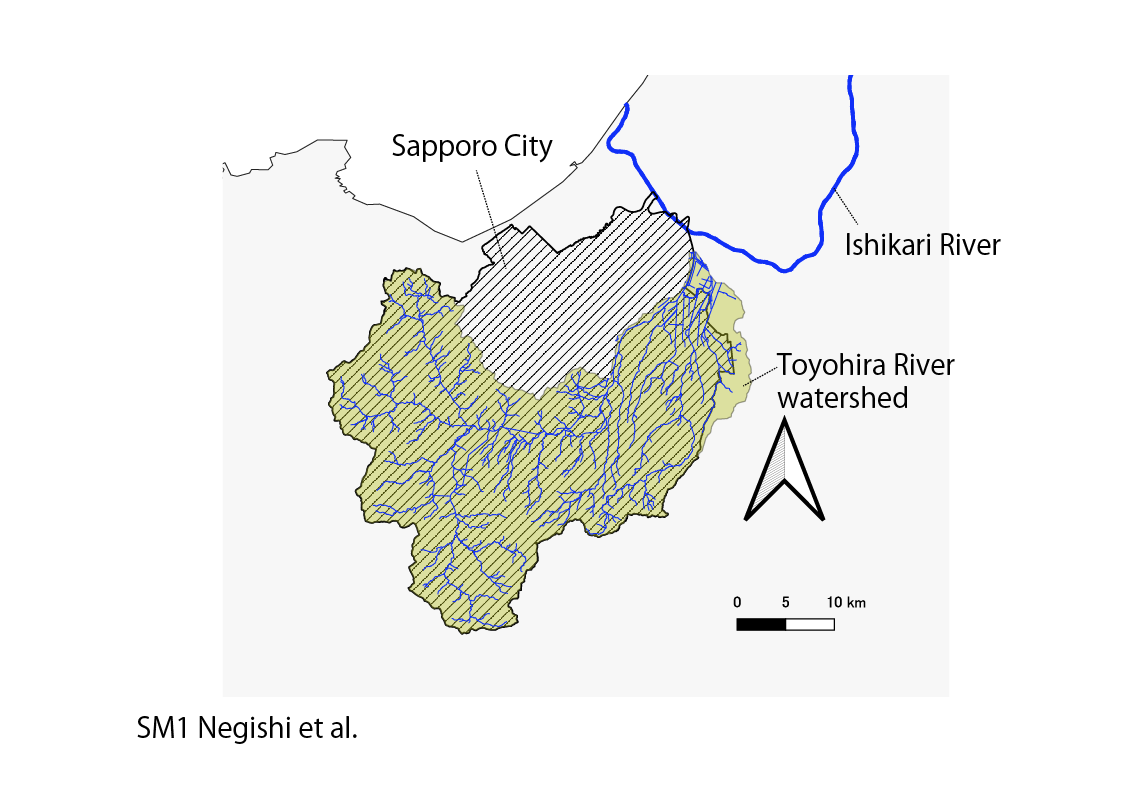

The area of Toyohira River watershed relative to the area of Sapporo City.

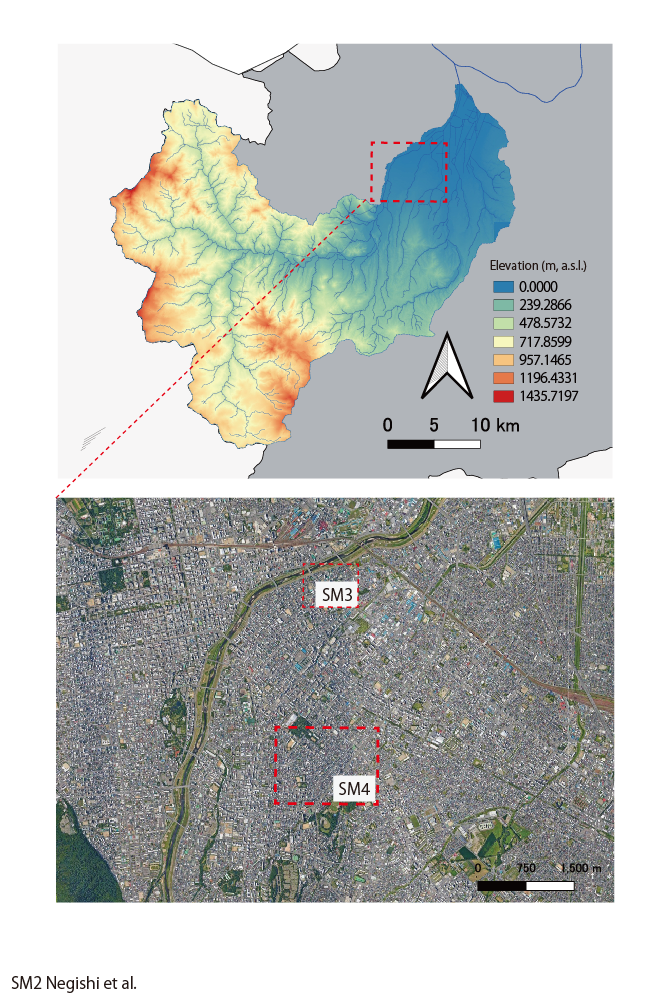

The lower panel shows the current condition of highly populated study area in Sapporo City; red dotted squares each denotes the area where temporal changes in landscape features were shown in details in SM3 and SM4.

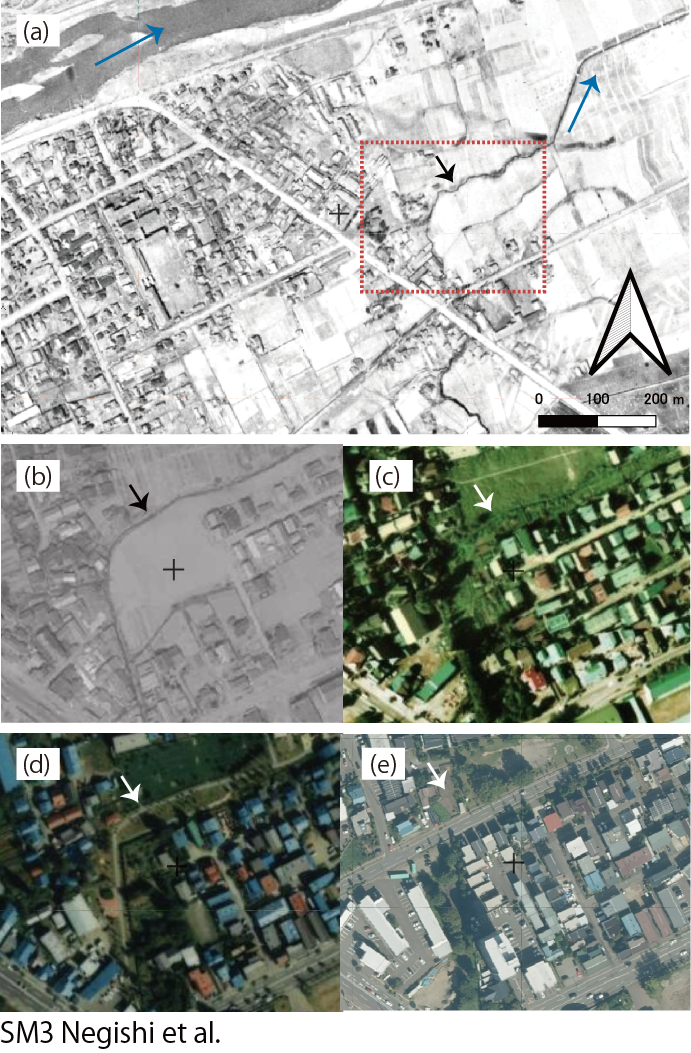

The landscape transformation over time from 1940s (a), 1960s (b), 1970s (c), 1980s (d), and 2020 (e). The area delineated by a red dotted square in (a) was shown in other panels. The allow in each panel denotes the exact same spot. Note that a river channel in (a) gradually disappeared over the time was completely replaced with other surface features.

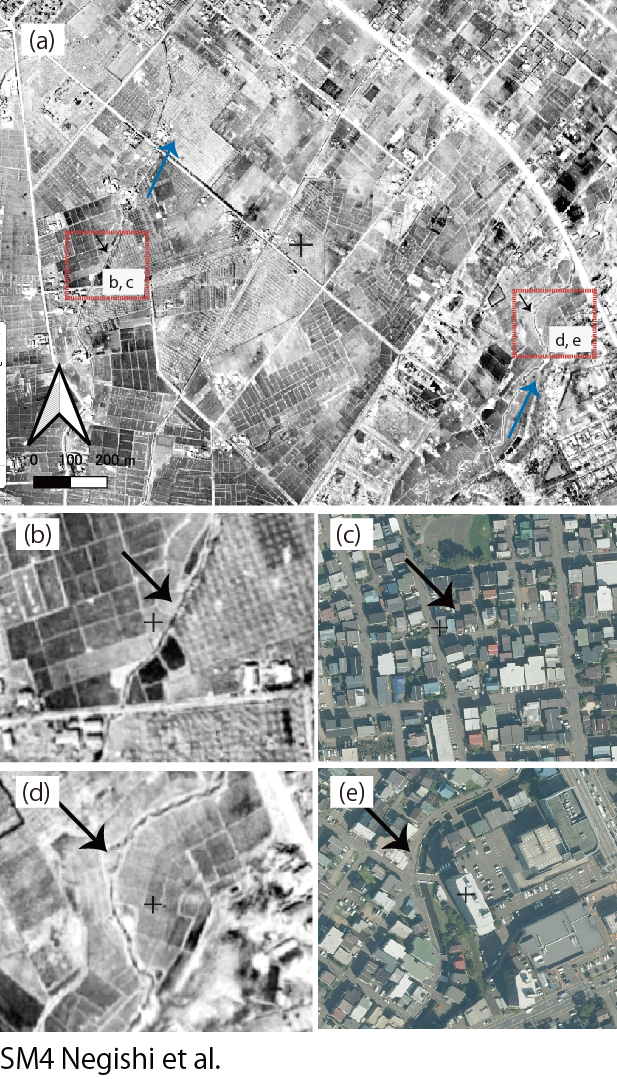

The landscape transformation over time from 1940s (b, d) to 2020 (c, e) for the areas each delineated by red dotted squares in (a). The allow in each panel denotes the exact same spot. Note that a river channel in (b) disappeared in (c) whereas a river channel in (d) exists in (e).

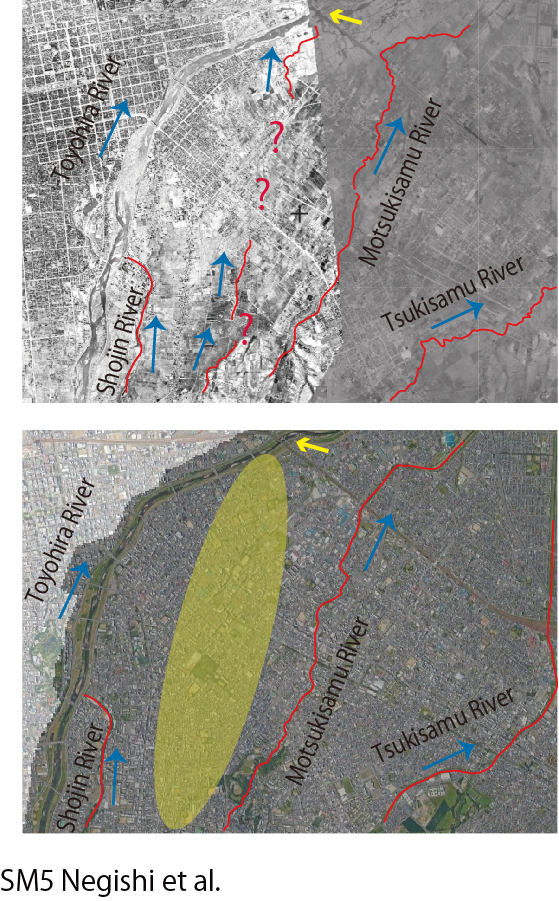

The upper panel shows landscape in 1940s to 1960s whereas the lower panel shows landscape in 2020. Yellow arrow indicates the location of the study segment. Question marks in the upper panels indicate the areas where river channel was not visually identified from the given air photos but was assumed to be present. Note that this tributary between Motsukisamu River and Shojin River was not visually identified or not confirmed by direct field observations on foot in 2020 (JNN, personal observations).

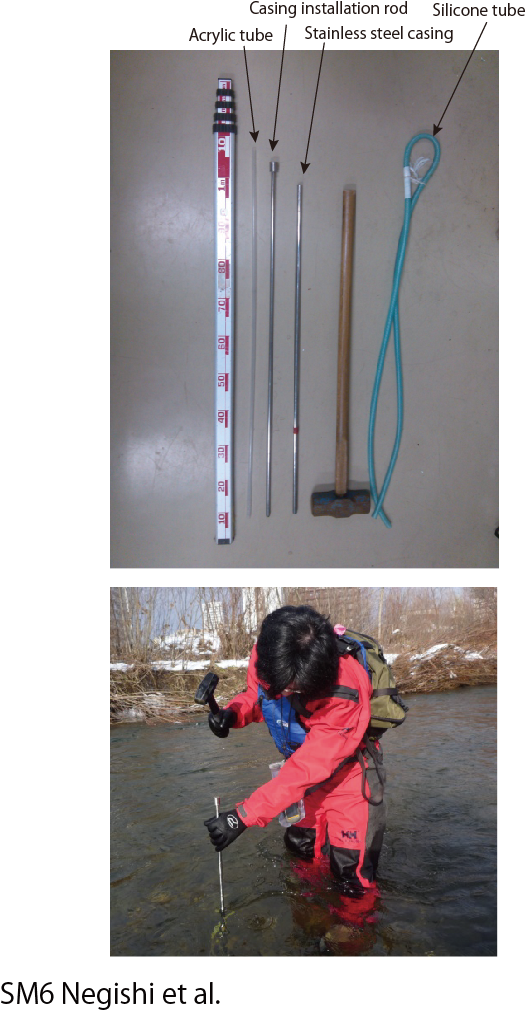

The upper panel shows mini-piezometer system used to collect hyporheic water. After stainless-steel casing (1 cm internal diameter) was pounded into 20-30 cm depth into the riverbed by using an installation rod, an acrylic tube (0.7 cm internal diameter) that was 1 m long, slotted along the bottom 5 cm, and capped at the end with 100-µm nylon mesh was inserted inside the casing. Then, casing was removed gently, after which silicone tube was connected to the top end of acrylic tube to collect water from the riverbed. At the spot of hyporheic water collection, surface water was also collected and in-situ measured for water quality parameters. The lower panel shows a photo that captured how casing installation was carried out.

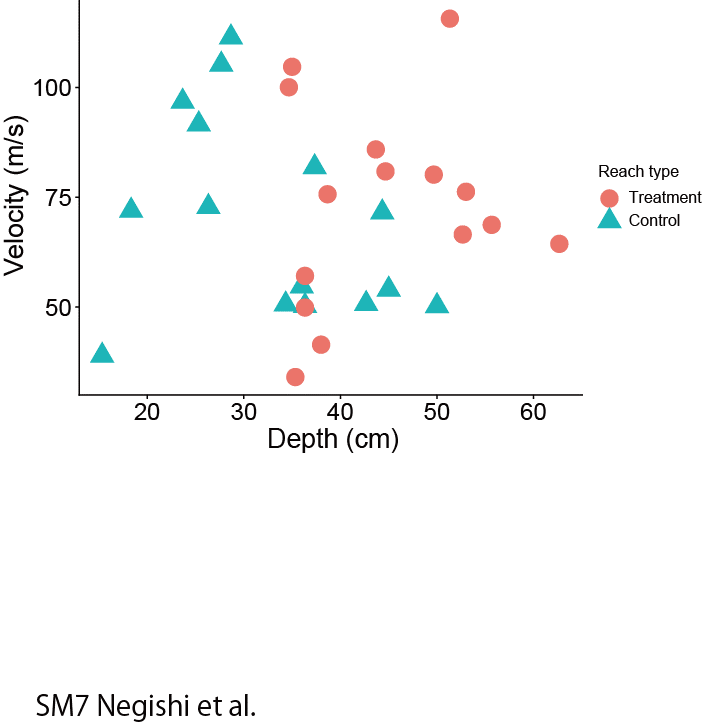

Water depth and velocity comparisons between two reaches. At each sampling points within each reach (a total of 15 points in respective reaches), three replicated measurements of water depth (cm) and flow velocity were made at the base-flow condition on August 8, 2016. Flow velocity was measured at 60% water depth using a propeller-based meter (model CR-11; Cosmo-riken Inc., Tokyo). Measurement values were averaged to represent data for each sampling point. Data were statistically compared between two reaches using PERMANOVA with Euclidian distance method, and the difference was insignificant (p=0.18).

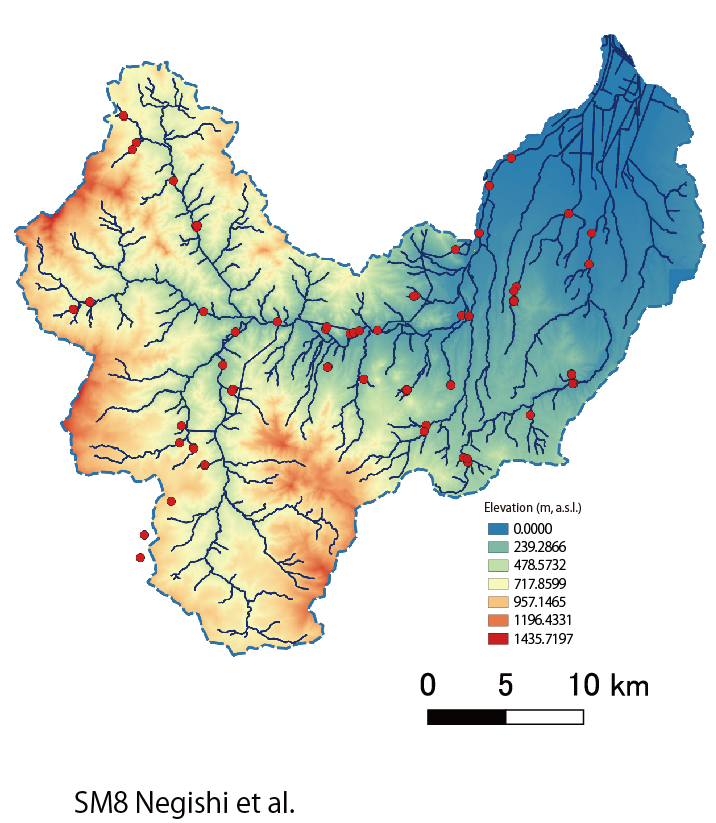

Red points denote locations of water collection for SIRs of hydrogen/oxygen in surface river water. Two points near the western upper boundary of catchment were outside of Toyohira River catchment because similarly high elevation sites could not be visited due to the absence of road accesses; samples collected from those sites were included as a part of Toyohira River watershed samples based on the assumption that they represent sample characteristics of sites within Toyohira River watershed having similar elevations.
